# Effects of phosphorylation on Drp1 activation by its receptors, actin, and cardiolipin

**DOI:** 10.1101/2023.08.20.554022

**Authors:** Ao Liu, Anna L. Hatch, Henry N. Higgs

**Affiliations:** Department of Biochemistry and Cell Biology, Geisel School of Medicine at Dartmouth, Hanover NH 03755

**Keywords:** K38A, CDK, PKA, S616, S637

## Abstract

Drp1 is a dynamin family GTPase that is required for mitochondrial and peroxisomal division, in which it oligomerizes into a ring and constricts the underlying membrane in a GTP hydrolysis-dependent manner. Oligomerization increases Drp1 GTPase activity through interactions between neighboring GTPase domains. In cells, Drp1 is regulated by several factors including Drp1 receptors, actin filaments, cardiolipin, and phosphorylation at two sites: S579 and S600. Phosphorylation of S579 is widely regarded as activating, while S600 phosphorylation is commonly considered inhibiting. However, the direct effects of phosphorylation on Drp1 GTPase activity have not been investigated in detail. In this study, we compare the effects of S579 and S600 phosphorylation on purified Drp1, using phospho-mimetic mutants and *in vitro* phosphorylation. The oligomerization state of both phospho-mimetic mutants is shifted toward smaller oligomers. Both phospho-mimetic mutations maintain basal GTPase activity, but eliminate GTPase stimulation by actin and decrease GTPase stimulation by cardiolipin, Mff, and MiD49. Phosphorylation of S579 by Erk2 produces similar effects. When mixed with wild-type Drp1, both S579D and S600D phospho-mimetic mutants reduce the actin-stimulated GTPase activity of Drp1-WT. Conversely, a Drp1 mutant that lacks GTPase activity, the K38A mutant, stimulates Drp1-WT GTPase activity under both basal and actin-stimulated conditions, similar to previous results for dynamin-1. These results suggest that the effect of S579 phosphorylation is not to activate Drp1 directly, and likely requires additional factors for stimulation of mitochondrial fission in cells. In addition, our results suggest that nearest neighbor interactions within the Drp1 oligomer affect catalytic activity.

## Introduction

The dynamin family GTPase Drp1 is an important mediator of membrane fission for at least two organelles, mitochondria and peroxisomes ^1^. When not mediating membrane fission, Drp1 is a cytoplasmic protein. To induce membrane fission, Drp1 is recruited to the target membrane by receptor proteins, where it oligomerizes into a ring structure around the membrane. Oligomerization increases Drp1 GTPase activity by bringing the N-terminal GTPase domains into close proximity ^2–4^. GTP hydrolysis results in constriction of the Drp1 ring and membrane constriction.

Drp1 receptors can directly activate Drp1. One Drp1 receptor, Mff, is on both mitochondrial and peroxisomal membranes. Mff itself is a trimer, and its binding causes increased Drp1 GTPase activity *in vitro* ^5,6^. A second set of Drp1 receptors, MiD49 and MiD51, are only found on mitochondria. In their inactive state, MiD proteins are monomeric and do not activate Drp1. Binding to fatty acyl-coenzyme A stimulates MiD oligomerization, which in turn stimulates Drp1 GTPase activity ^7^.

Other regulatory molecules for Drp1 include actin filaments and phospholipids. Biochemically, actin filaments stimulate Drp1 GTPase activity ∼4-fold through direct binding ^8,9^. In cells, actin polymerization through an endoplasmic reticulum-bound formin protein, INF2, increases Drp1 oligomerization and mitochondrial recruitment ^8,10,11^. The mitochondrial lipid cardiolipin (CL) can affect a >20-fold stimulation of GTPase activity biochemically ^12^, while CL-derived phosphatidic acid suppresses Drp1 activation ^13^.

Drp1 is also subject to a number of post-translational modifications ^14,15^, with phosphorylation in particular being correlated with changes in Drp1-mediated mitochondrial fission. Two well-studied phosphorylation sites occur within a 21 amino acid segment of the Variable Domain of Drp1^15^ (**Figure 1A),** which forms an unstructured loop at one end of the elongated Drp1 structure^2^ (**Figure 1B**). Depending on the splice variant and species studied ^16,17^, the positions of these sites differ (**Figure 1C**), with the most common names in the literature being S616 and S637, but which will be referred to here mostly as S579 and S600, as explained in the Results section.

**Figure 1.**
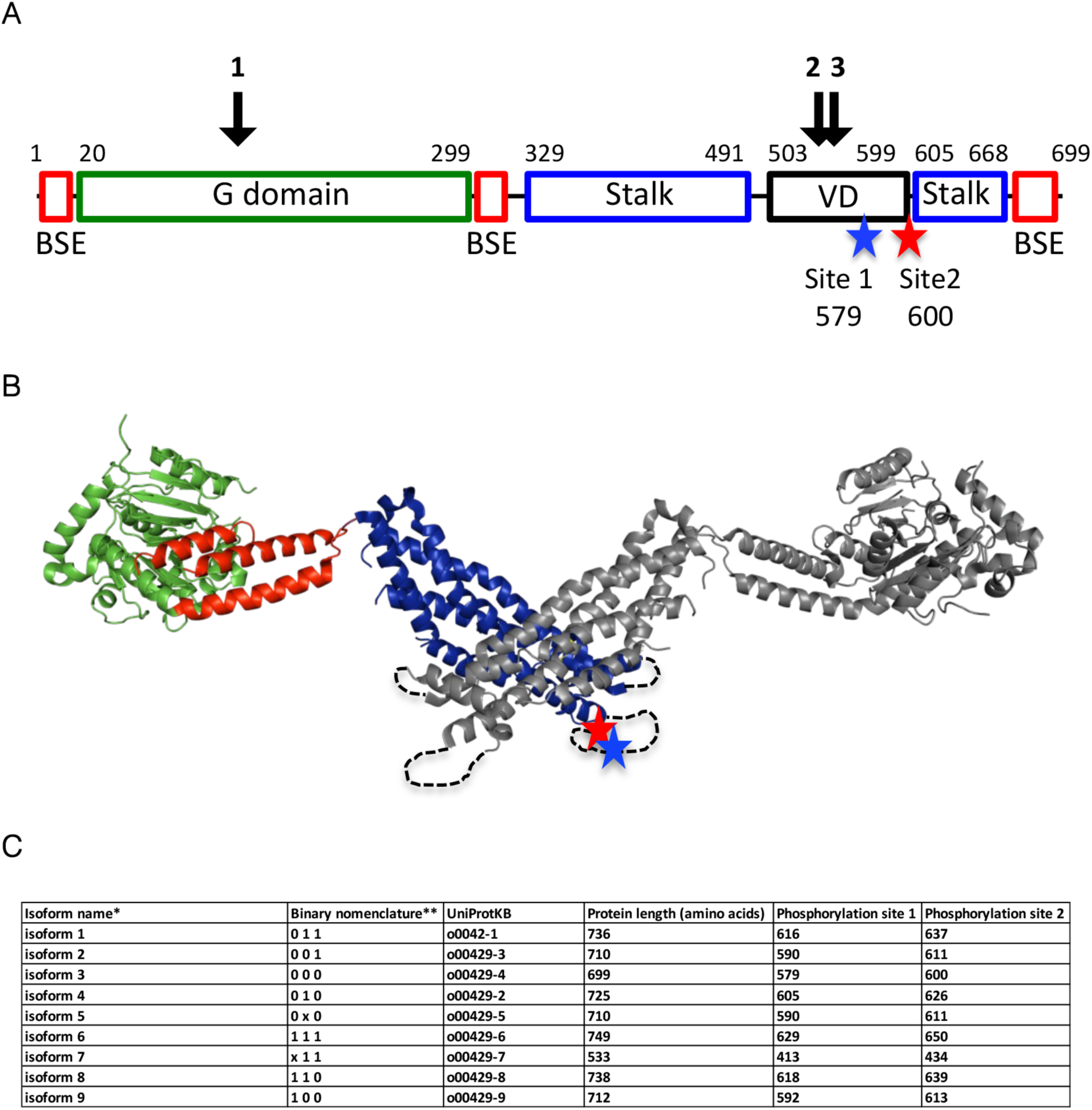
Drp1 phosphorylation sites. (A) Domain organization of Drp1 isoform 3 (also called Drp1-000): GTPase domain (green), Bundle Signaling Element (BSE, red), stalk (blue), Variable Domain (VD, black, also called the B-insert in some publications). Blue and red stars indicate phosphorylation sites S579 (site 1) and S600 (site 2), respectively. Black arrows indicate locations of the three alternatively spliced inserts. (B) Structural model of Drp1 dimer (PDB 4BEJ) showing positions of phosphorylation sites S579 (blue star) and S600 (red star) on one subunit (color coded similar to panel A). Dashed loops for Variable Domain denote that this was not resolved in PDB 4BEJ. (C) Table listing positions of the phosphorylation sites corresponding to S579 (Site 1) and S600 (Site 2) in this paper for the nine Drp1 isoforms listed in UniProt, following isoform designation given by ^17^ and binary nomenclature used by ^16^. Binary nomenclature based on presence (1) or absence (0) of the three alternatively spliced inserts, with ‘x’ denoting a variation of the indicated site (described in ^17^).

Phosphorylation of S579 has been observed for multiple kinases, including by CDK1, CDK5, ERK2, CaMKII, ROCK, PKCδ, and PINK ^16,18–25^, with phosphorylation being correlated with increased mitochondrial fission. Phosphorylation of S600 by protein kinase A (PKA), CaMK1a, ROCK1, AMPK and protein kinase D has been reported ^26–31^. In most cases, S600 phosphorylation has been correlated with decreased mitochondrial fission^26,27,30,32^. However, some studies show evidence for a positive effect of S600 phosphorylation on cellular Drp1 activity ^28,29,31,33^, while another study reports no effect ^34^. Finally, one report suggests that S600 phosphorylation promotes S579 phosphorylation, and that the doubly phosphorylated S579/S600 protein correlates with increased mitochondrial fission ^35^. Many of the above-cited publications utilize phospho-mimetic mutants to induce similar mitochondrial changes to phosphorylation-induced changes, suggesting that phospho-mimetics can elicit similar effects.

It is unclear whether phosphorylation at either site directly alters Drp1 activity. For S579 phosphorylation, no biochemical data addressing Drp1 activity are available. For S600 phosphorylation, the biochemical data are conflicting. In one study, a phospho-mimetic mutant of Drp1 maintains oligomerization and GTP hydrolysis activity ^27^. In another study, *in vitro* phosphorylation of GST-Drp1 by PKA inhibits GTPase activity ^26^. Recent structural work suggests that S600 phosphorylation inhibits MiD49 binding^36^, although other cellular co-IP experiments suggest that the S600D phospho-mimetic binds better to MiD49 and MiD51 than does the non-phosphorylatable S600A mutant ^37^. Other studies show that the region of Drp1 containing both phosphorylation sites (the Variable Domain) inhibits binding to Mff ^5,38^, although neither S579D nor S600D phospho-mimetics alter this effect ^38^.

In this study, we assess the biochemical effects of Drp1 phosphorylation on oligomerization and GTPase activity of Drp1 alone, and in the presence of several activators (actin, cardiolipin, Mff, MiD49). For these studies, we use both phospho-mimetic S-to-D mutations and *in vitro* phosphorylation on the S579 site. Surprisingly, we find that both types of phosphorylation inhibit the abilities of these activators to stimulate Drp1 activity.

## Results

### Phospho-mimetic mutants are less oligomerized in the nucleotide-free state

We constructed two phospho-mimetic Drp1 mutants corresponding to S616D and S637D in isoform 1. We refer to these mutants, however, as S579D and S600D respectively, because we use isoform 3 of Drp1 (**Figure 1C**). In HeLa, HL60 and PC12 cells, isoform 3 is the most abundant isoform present, making up over 40% of total Drp1 protein ^16^.

The S579 site, generally thought to activate Drp1, is located within the Variable Domain; while the S600 site, generally thought to inhibit Drp1, marks the boundary between the Variable Domain and the stalk (**Figure 1A**). The region encompassing both sites is poorly resolved in the existing structural models of nucleotide-free Drp1^2^ (**Figure 1B**) or other models ^36^.

By size exclusion chromatography at high Drp1 concentration (30 μM), both phospho-mutants are slightly shifted to smaller sizes compared to Drp1-WT (**Figure 2A**). For more detailed analysis of Drp1 hydrodynamic properties, we used velocity analytical ultracentrifugation (vAUC). Previously, we reported that purified Drp1-WT exists in several oligomeric states ^9^, similar to the results of other studies^2,12^. We compared the Drp1-S579D and Drp1-S600D to Drp1-WT by vAUC at three concentrations: 8, 4, and 1.5 µM (**Figure 2B**). As with Drp1-WT, both mutants display a 7 S species that is similar to the sedimentation pattern of a oligomerization-deficient mutant ^2,9^, suggesting that this is the dimeric species. In addition, several species of higher S values, corresponding to larger oligomers, are present for Drp1-WT. At all concentrations tested, both Drp1 S579D and S600D shift towards smaller oligomers when compared with WT, with the S600D mutant displaying a greater shift.

**Figure 2.**
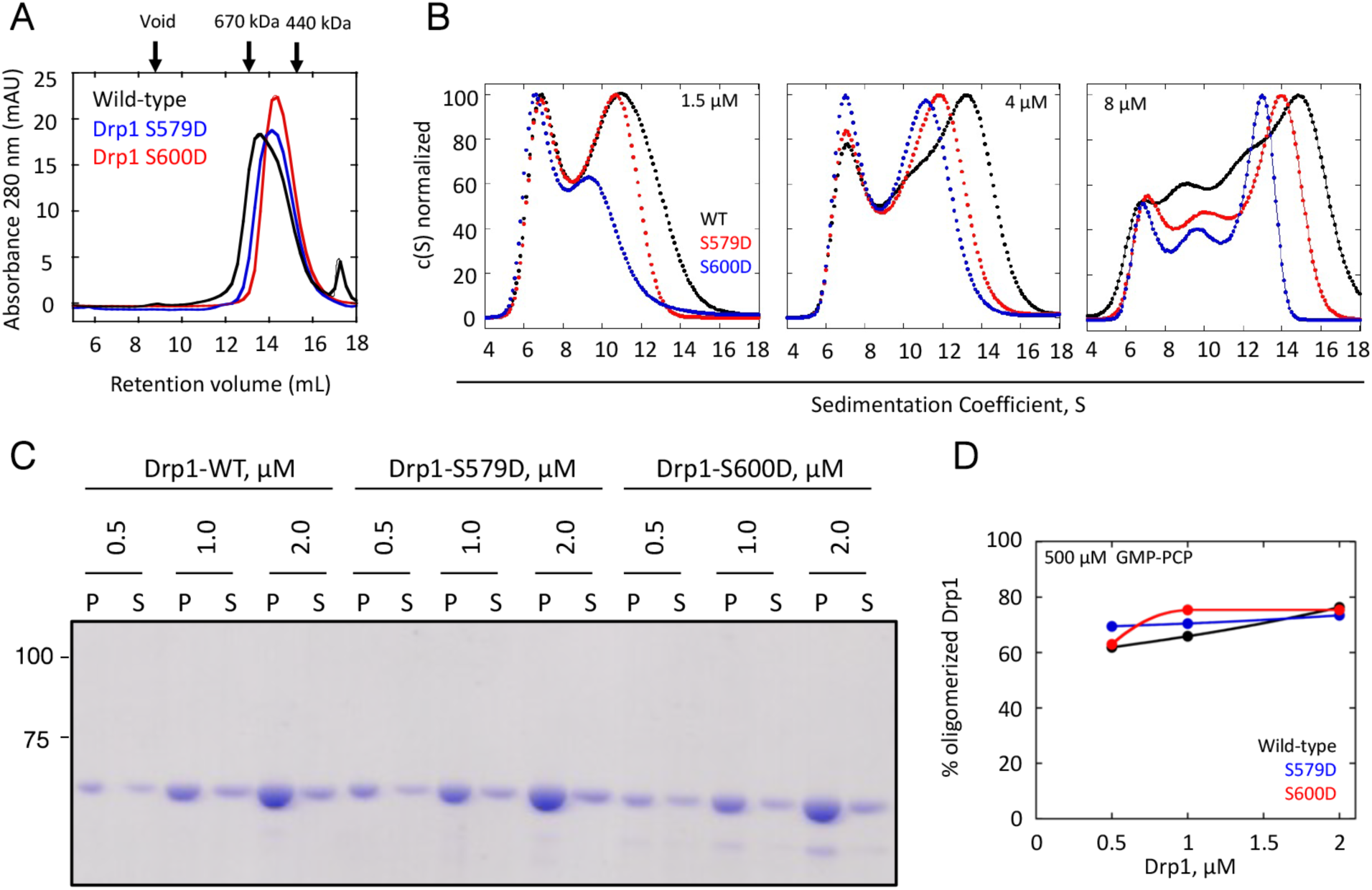
Oligomeric properties of Drp1 phospho-mimetic mutants. (A) Superose 6 gel filtration profiles of 30 µM of Drp1-WT, Drp1-S579D, and Drp1-S600D. At top are peak elution positions for two standards (thyroglobulin (670 kDa) and ferritin (440 kDa)) as well as the void volume position. (B) Velocity analytical ultracentrifugation of Drp1-WT, Drp1-S579D, and Drp1-S600D (black, red and blue respectively) at three concentrations: 8, 4 and 1.5 µM. (C) Coomassie-stained gel showing sedimentation assays graphed in panel C. Mass markers (in kDa) shown at left. P = pellet, S = supernatant. (D) Quantification of % oligomerized Drp1 from sedimentation assays of Drp1-WT, Drp1-S579D, and Drp1-S600D in the presence of the non-hydrolysable GTP analogue GMP-PCP (500 µM) at 0.5, 1.0, and 2.0 µM Drp1.

We also tested the ability of Drp1 to form larger oligomers in the presence of the non-hydrolysable GTP analogue GMP-PCP using a high-speed sedimentation assay^9^. At three Drp1 concentrations (0.5, 1.0, and 2.0 µM), GMP-PCP causes similar degrees of Drp1 sedimentation for all constructs, suggesting that GTP-mediated oligomerization is not affected by these mutations (**Figure 2 C,D**).

These data suggest that both the S579D and S600D mutants display less ability to oligomerize in the nucleotide-free state than WT Drp1, but that non-hydrolyzable GTP induces all three proteins to oligomerize.

### Phospho-mimetic mutants for both S579 and S600 reduce Drp1 responses to activators

We next examined the GTPase activities of the two phospho-mimetic mutants. Both mutants display similar GTPase activity to Drp1-WT (1.57±0.09, 1.51±0.02, and 1.33±0.14 μM/min/μM for WT, S579D, and S600D, respectively). Previously, we reported that actin filaments have a bi-phasic effect on Drp1 GTPase activity, where low actin filament concentrations are stimulatory but higher concentrations bring the activity back to the Drp1-alone values ^9^. We tested whether the phospho-mimetic mutants responded to actin in a similar manner. Surprisingly, actin does not activate Drp1 S579D or Drp1 S600D at any concentration tested (**Figure 3A**).

**Figure 3.**
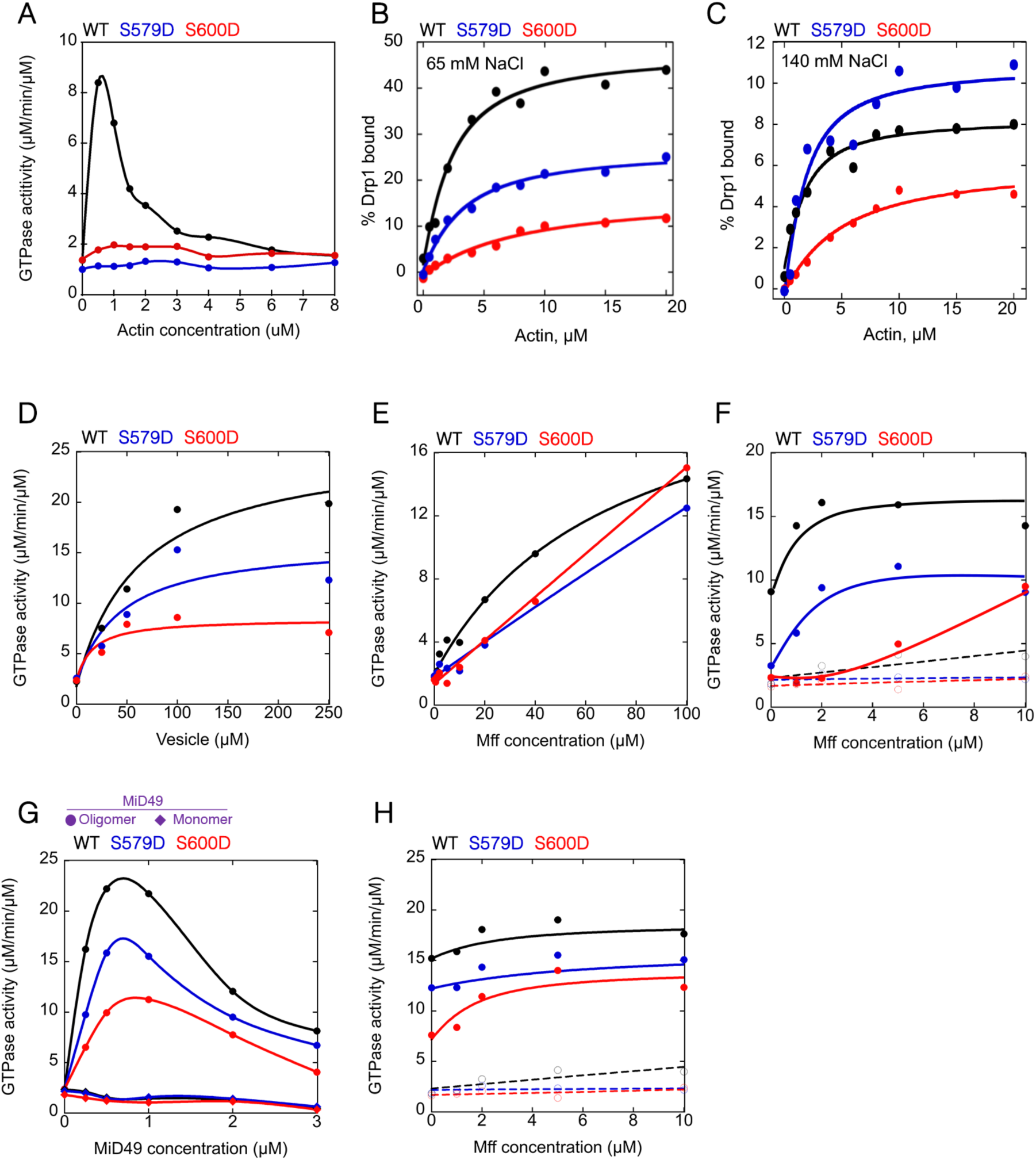
Effects of Drp1 phospho-mimetic mutants on actin binding and GTPase stimulation by actin, Mff, and MiD49. Wild-type, Drp1 S579D, Drp1 S600D shown in black, blue, and red respectively for all panels. All GTPase assays contain 0.75 µM Drp1. (A) GTPase assays containing 0.75 µM Drp1 that was pre-incubated with the indicated concentration of actin filaments for 5 min before GTP addition. Activity expressed as µM phosphate released per minute per µM Drp1. (B) and (C) Graphs of % Drp1 bound versus actin concentration, from co-sedimentation assays at 65 mM (B) and 140 mM NaCl (C), respectively (1.3 µM Drp1 in all cases). Raw data in Supplemental Figure 1. (D) GTPase assays containing Drp1 that was pre-incubated with the indicated concentration of cardiolipin-containing vesicles (40% DOPC, 35% DOPE, 25% CL) for 5 min before GTP addition. (E) Comparison of Drp1-WT with Drp1-S579D and Drp1-S600D GTPase activities in the presence of the indicated concentration of Mff for 5 min before GTP addition. (F) Comparison of Drp1-WT with Drp1-S579D and Drp1-S600D GTPase activities in the presence of 0.5 µM actin filaments and the indicated concentration of Mff for 5 min before GTP addition. Values for Mff alone are indicated by open circles and dashed lines, while values for Mff/actin are indicated by closed circles and solid lines. (G) Comparison of Drp1-WT with Drp1-S579D and Drp1-S600D GTPase activities in the presence of the indicated concentration of MiD49 monomer (diamonds) or MiD49 oligomer (circles) for 5 min before GTP addition. (H) Comparison of Drp1-WT with Drp1-S579D and Drp1-S600D GTPase activities in the presence of 0.25 µM MiD49 oligomer and the indicated concentration of Mff for 5 min before GTP addition. Values for Mff alone are indicated by open circles and dashed lines, while values for Mff/MiD49 are indicated by closed circles and solid lines.

Previously, we reported that Drp1 binds actin filaments with a Kd^app^ in the range of 1-2 µM ^8,9^. We compared actin binding for Drp1-WT, Drp1-S579D and Drp1-S600D by co-sedimentation assay at two ionic strengths: 65 mM and 140 mM NaCl. Drp1-WT and Drp1-S579D display similar Kd for actin at 65 mM NaCl (Kd^app^ 1.7 and 2.5 µM for WT and S579D, respectively), while Drp1-S600D displays significantly lower actin affinity (Kd^app^ 7.1 µM). We previously reported that Drp1-WT binding to actin filaments saturates at approximately 50% Drp1 bound ^9^. Both mutants display lower % bound at saturating actin (48%, 27%, 18% bound for WT, S579D, and S600D respectively) (**Figure 3B**; **Supplemental Figure S1A, B**). At 140 mM NaCl, Drp1 S579D has a similar affinity for actin filaments as WT (1.1 and 1.2 µM for S579D and wild-type respectively) and a comparable % bound (10.9% and 8.3% for S579D and WT respectively), while Drp1 S600 has lower affinity and % bound than the other constructs (4.3 µM and 6.1% bound) (**Figure 3C; Supplemental Figure S1C, D, E**). These results show that the phospho-site mutants have altered actin-binding properties, with Drp1 S600D displaying less actin interaction under all conditions.

We also asked whether GTPase stimulation by another Drp1 activator, cardiolipin (CL) ^12^, was affected by the S579D or S600D mutations. Interestingly, unilamellar vesicles containing 25 mole% cardiolipin are less stimulatory to Drp1-S579D and Drp1-S600D than they are for Drp1-WT (**Figure 3D**).

We have previously shown that the cytoplasmic region of the Drp1 receptor Mff activates Drp1, and that actin filaments synergize with Mff by reducing the Mff concentration needed for maximal Drp1 activation ^6^. Both phospho-mimetic mutants display slightly decreased activation by Mff alone (**Figure 3E**). Interestingly, actin filaments still synergize with Mff for Drp1-S579D stimulation, albeit with a lower maximal activation (**Figure 3F**). In contrast, the Drp1-S600D mutant displays greatly reduced synergy between actin and Mff (**Figure 3F**).

Two other Drp1 receptors, MiD49 and MiD51, do not activate Drp1 GTPase activity when they themselves are not activated. Upon binding their activating ligand, acyl-CoA, MiD proteins oligomerize, resulting in Drp1 GTPase activation in a bi-phasic manner, similar to actin ^7^. Here, we find that both Drp1 phospho-mimetic mutants are also activated by acyl-CoA-bound MiD49 in a bi-phasic manner. However, the degree of activation is lower, with Drp1-S600D being most affected (**Figure 3G**). Similar to Drp1-WT, neither phospho-mimetic mutant is stimulated by MiD49 monomer (**Figure 3G**).

As with actin filaments, oligomerized MiD49 synergizes with Mff by reducing the Mff concentration required for maximal Drp1 activation. Drp1-S579D displays maximal activity in the presence of low concentrations of Mff and oligomerized MiD49, albeit at a lower maximum than Drp1-WT (**Figure 3H**). Similarly, despite its decreased ability to stimulate Drp1-S600D, oligomerized MiD49 is able to reduce the concentration of Mff required for Drp1-S600D stimulation.

These results show that phospho-mimetics of both the ‘activating’ (S579D) and ‘inhibitory’ (S600D) sites on Drp1 result in a reduction of Drp1 activation by a variety of activators. This reduction is most dramatic for actin filaments.

### Drp1 phosphorylated on S579 displays similar properties to the S579D phospho-mimetic

The results for Drp1-S579D were surprising, considering that this site has been found to be associated with increased mitochondrial fission in cells. We sought to test these results further by directly phosphorylating Drp1 on S579 using recombinant ERK2, one of the kinases shown to mediate this phosphorylation ^20,21^ Incubation with ERK2 for 4-hr results in a slight decrease in mobility on SDS-PAGE, suggestive of phosphorylation (**Figure 4A**). Western blotting using an antibody against phospho-S579 shows saturation of this signal on a similar time scale (**Figure 4B**). Gel filtration chromatography of 4-hr phosphorylated Drp1 results in a similar elution profile to mock-phosphorylated sample (**Figure 4C**, 30 µM Drp1 loaded). Analysis by tryptic digest and mass spectrometry reveals that phospho-S579 accounts for 94.5% of the phosphorylated peptides identified, with other detectable phosphorylation sites being S526 (5.2%), S136 (0.3%), S570 (0.02%), and T394 (0.01%). No phosphorylated peptides were detected for mock-phosphorylated Drp1. These results suggest that our ERK2-treated Drp1 is efficiently phosphorylated on S579, with minor phosphorylation on other residues.

**Figure 4.**
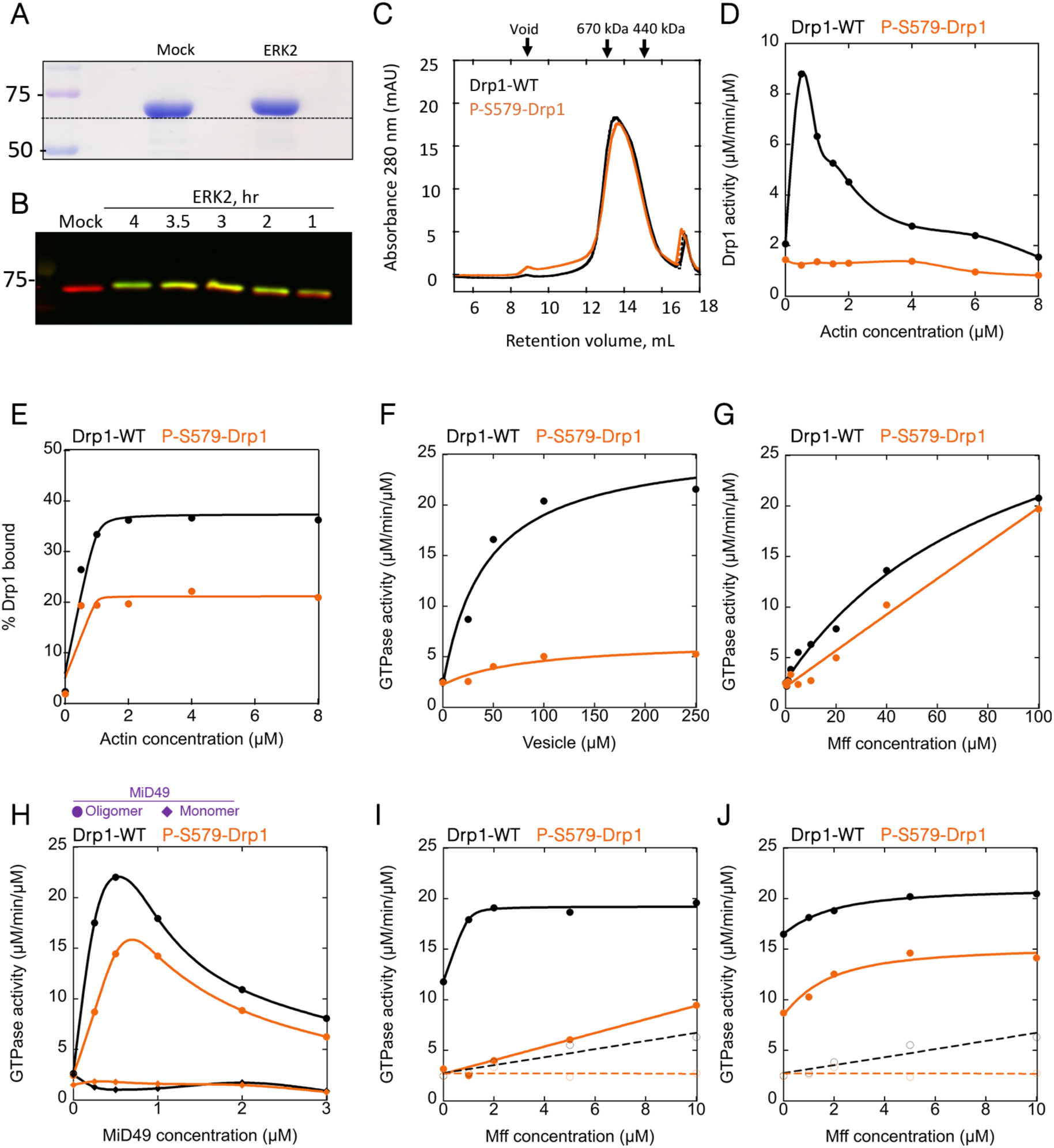
Effects of Phospho-S579-Drp1 on actin binding and GTPase stimulation by actin, Mff, and MiD49. In all graphs, Drp1-WT and P-S579-Drp1 in black and orange, respectively. All GTPase assays contain 0.75 µM Drp1. (A) Coomassie-stained SDS-PAGE of Drp1 from either mock phosphorylation or ERK2 phosphorylation reactions (4 hr). 2 µg Drp1 on gel. Positions of size standards (in kDa) on left. (B) Western blot showing anti-Drp1 (red) and anti-phospho-S579-Drp1 (green) at varying times after ERK2 treatment. (C) Superose 6 gel filtration profiles of WT and P-S579-Drp1. At top are peak elution positions for two standards (thyroglobulin (670 kDa) and ferritin (440 kDa)) as well as the void volume position. (D) GTPase assays containing Drp1 that was pre-incubated with the indicated concentration of actin filaments for 5 min before GTP addition. Activity expressed as µM phosphate released per minute per µM Drp1. (E) Graph of % Drp1 bound versus actin concentration, from co-sedimentation assays at 65 mM NaCl. Raw data in Supplemental Figure S2. (F) GTPase assays containing Drp1 that was pre-incubated with the indicated concentration of cardiolipin-containing vesicles (μM total lipid, vesicles contain 40% DOPC, 35% DOPE, 25% CL) for 5 min before GTP addition. (G) Comparison of WT and P-S579-Drp1 GTPase activities in the presence of the indicated concentration of Mff for 5 min before GTP addition. (H) Comparison of WT and P-S579-Drp1 GTPase activities in the presence of the indicated concentration of MiD49 monomer (diamond) or MiD49 oligomer (circle) for 5 min before GTP addition. (I) Comparison of WT and P-S579-Drp1 GTPase activities in the presence of 0.5 µM actin filaments and the indicated concentration of Mff for 5 min before GTP addition. Values for Mff alone are indicated by open circles and dashed lines, while values for Mff/actin are indicated by closed circles and solid lines. (J) Comparison of WT and P-S579-Drp1 GTPase activities in the presence of 0.25 µM MiD49 oligomer and the indicated concentration of Mff for 5 min before GTP addition. Values for Mff alone are indicated by open circles and dashed lines, while values for Mff/MiD49 are indicated by closed circles and solid lines.

Similar to its phospho-mimetic analogue, S579-phosphorylated Drp1 (P-S579-Drp1) is not stimulated by actin filaments (**Figure 4D**). In actin binding assays, P-S579-Drp1 displays a ∼2-fold decrease in maximal actin binding relative to Drp1-mock, while maintaining a similar affinity (**Figure 4E, Supplementary Figure S2**), similar to the S579D mutant. Cardiolipin-containing vesicles only weakly stimulate P-S579-Drp1 (**Figure 4F**), which is a comparatively greater effect than that of the phospho-mimetic. Similar to Drp1-S579D, P-S579-Drp1 is stimulated by Mff or by MiD49 oligomers to a lesser extent than Drp1-WT (**Figure 4G,H**). Interestingly, the synergy of Mff with actin filaments is strongly reduced for Drp1-S579D (**Figure 4I**) while the synergy between Mff and MiD49 oligomers is maintained (**Figure 4J**).

A recent publication reports that phosphorylation at the S600 position can stimulate S579 phosphorylation in some cases, and this doubly phosphorylated Drp1 leads to increased mitochondrial fragmentation ^35^. We therefore tested whether doubly phosphorylated Drp1 displays different properties to either singly phosphorylated protein. For these experiments, we phosphorylated Drp1-S600D on S579 using Erk2, resulting in a similar gel shift as Drp1-phosphoS579 (**Figure 5A**). Similarly to the Erk2-phosphorylated Drp1-WT, Drp1-phosphoS579/S600D is not activated by actin filaments or cardiolipin (**Figure 5B,C**), and displays reduced activation by Mff (**Figure 5D**). Interestingly, Drp1-phosphoS579/S600D displays no activation by MiD49 oligomers (**Figure 5E**), and no synergistic effect of actin or MiD49 oligomers on Mff activation (**Figure 5F, G**).

**Figure 5.**
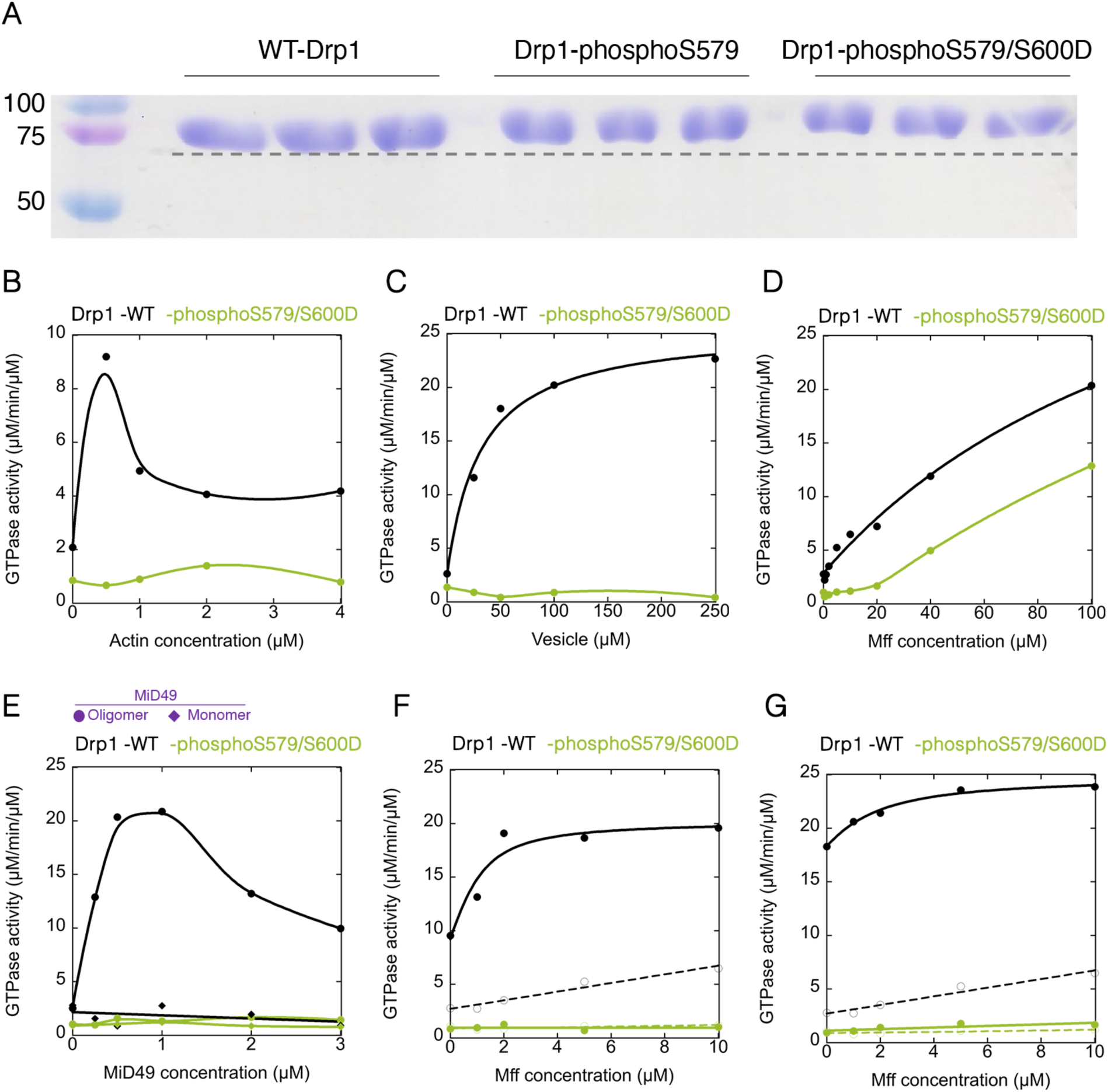
Effects of Drp1-phosphoS579/S600D on GTPase stimulation. In all graphs, Drp1-WT and Drp1-phosphoS579/S600D in black and green, respectively. All GTPase assays contain 0.75 µM Drp1. (A) Coomassie-stained SDS-PAGE of Drp1 from either Drp1-mock phosphorylation, Drp1-ERK2 phosphorylation, or Drp1-S600D-ERK2 phosphorylation reactions (4 hr). 2 µg Drp1 on gel. Positions of size standards (in kDa) on left. (B) GTPase assays containing Drp1 that was pre-incubated with the indicated concentration of actin filaments for 5 min before GTP addition. Activity expressed as µM phosphate released per minute per µM Drp1. (C) GTPase assays containing Drp1 that was pre-incubated with the indicated concentration of cardiolipin-containing vesicles (40% DOPC, 35% DOPE, 25% CL) for 5 min before GTP addition. (D) Comparison of Drp1-WT and Drp1-phosphoS579/S600D GTPase activities in the presence of the indicated concentration of Mff for 5 min before GTP addition. (E) Comparison of Drp1-WT and Drp1-phosphoS579/S600D activities in the presence of the indicated concentration of MiD49 monomer (diamond) or MiD49 oligomer (circle) for 5 min before GTP addition. (F) Comparison of Drp1-WT and Drp1-phosphoS579/S600D GTPase activities in the presence of 0.5 µM actin filaments and the indicated concentration of Mff for 5 min before GTP addition. Values for Mff alone are indicated by open circles and dashed lines, while values for Mff/actin are indicated by closed circles and solid lines. (G) Comparison of Drp1-WT and Drp1-phosphoS579/S600D activities in the presence of 0.25 µM MiD49 oligomer and the indicated concentration of Mff for 5 min before GTP addition. Values for Mff alone are indicated by open circles and dashed lines, while values for Mff/MiD49 are indicated by closed circles and solid lines.

The Drp1 protein used thus far (isoform 3) contains none of the three possible alternately spliced inserts. While isoform 3 predominates in some cell types, it is a minor isoform in others, and the other splice variants also make up an appreciable percentage of total Drp1 in most cells and tissues ^16^. We asked whether the isoform containing all three splice inserts (isoform 6) reacts similarly to isoform 3 in response to Erk2 phosphorylation. Treatment of isoform 6 with Erk2 results in decreased mobility on SDS-PAGE (**Figure 6A**), suggesting successful phosphorylation on S629 (equivalent to S579 in isoform 3). Similar to isoform 3, P-S629-Drp1(i6) is not activated by actin filaments (**Figure 6B**). In addition, Mff and MiD49 stimulate P-S629-Drp1(i6) to a lesser extent than non-phosphorylated Drp1 (**Figure 6 C,D**). These results suggest that the presence of the three splice inserts do not alter Drp1’s response to phosphorylation on the canonical ‘activating’ site.

**Figure 6.**
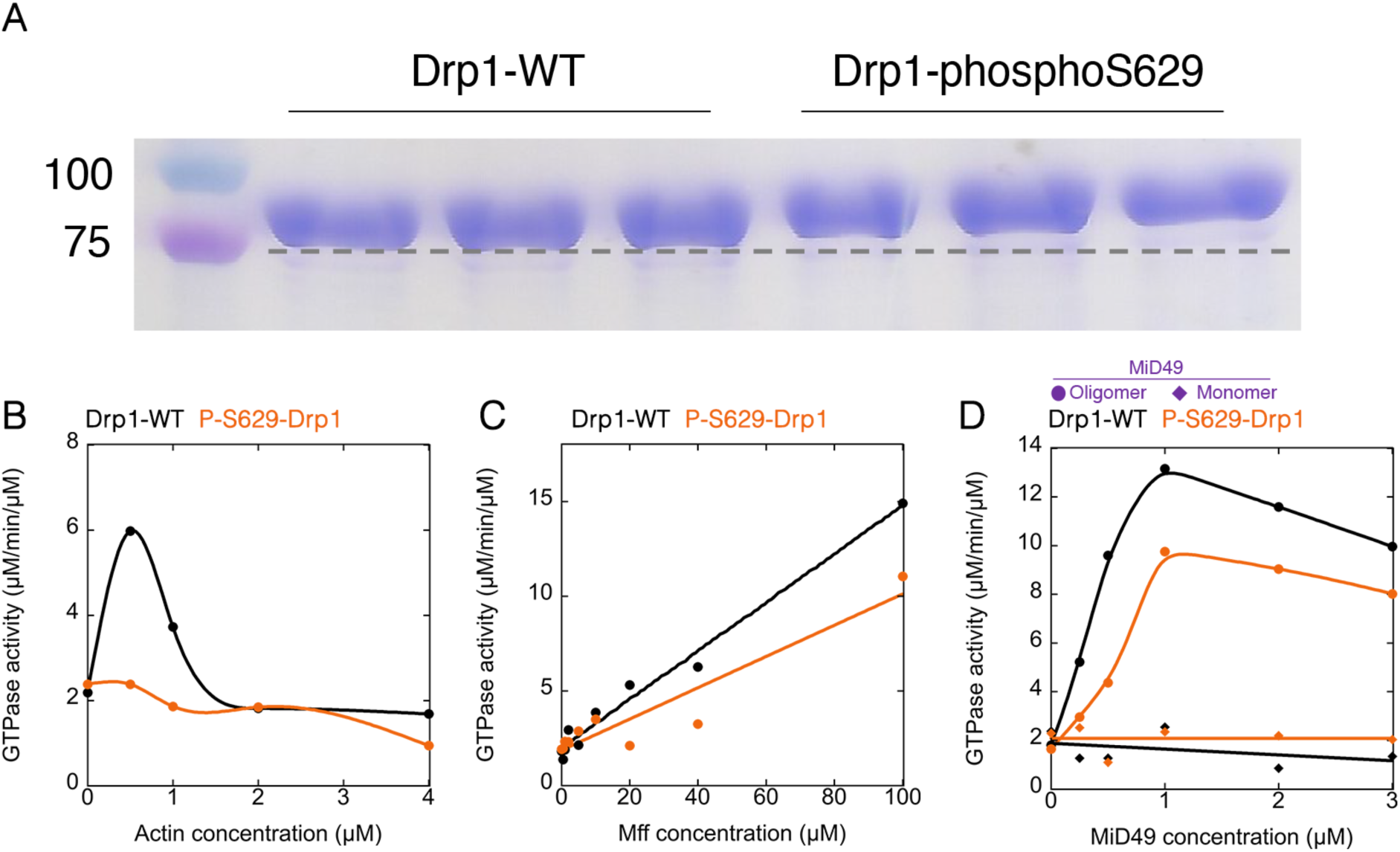
Effects of Erk2 phosphorylation on GTPase activity of Drp1 isoform 6. In all graphs, Drp1-WT and Drp1-phosphoS629 in black and orange, respectively. All GTPase assays contain 0.75 µM Drp1. (A) Coomassie-stained SDS-PAGE of Drp1 isoform 6 from either Drp1- mock phosphorylation or Drp1- ERK2 phosphorylation (4 hr). 2 µg Drp1 on gel. Positions of size standards (in kDa) on left. (B) GTPase assays containing Drp1 that was pre-incubated with the indicated concentration of actin filaments for 5 min before GTP addition. Activity expressed as µM phosphate released per minute per µM Drp1. (C) Comparison of Drp1-WT and P-S629-Drp1 GTPase activities in the presence of the indicated concentration of Mff for 5 min before GTP addition. (E) Comparison of Drp1-WT and P-S629-Drp1 activities in the presence of the indicated concentration of MiD49 monomer (diamonds) or MiD49 oligomer (circles) for 5 min before GTP addition.

These results show that Drp1-phosS579 possesses similar properties to the phospho-mimetic S579D mutant in its reduced stimulation by all tested activators. In fact, Drp1-phosS579 displays less stimulation by cardiolipin than Drp1-S579D, and displays no synergy between actin and Mff.

### Drp1-S579D does not stimulate actin-mediated activation of Drp1-WT

The ability of the S579D to bind actin without stimulation of its GTPase activities allowed us to test the effects of Drp1-S579D on actin-mediated activation of Drp1-WT GTPase activity. Our reasoning was as follows: if GTPase activation is a consequence of interaction between two neighboring GTPase domains, heteromeric assemblies between phosphorylated and non-phosphorylated Drp1 might have effects on GTPase activation. We used GFP-tagged Drp1 as the WT version of Drp1, to distinguish between WT and mutant proteins in co-sedimentation assays. In previous studies, we found that GFP-Drp1 also bound actin filaments^9^.

First, we varied percentages of Drp1-S579D and GFP-Drp1-WT, while maintaining a constant total Drp1 concentration, to assess effects on both actin binding and actin-stimulated GTPase activity by the WT construct. In these experiments, we used concentrations of total Drp1 (1.3 µM) and actin (1 µM) that result in sub-saturating Drp1 on actin. There is a linear increase in the % GFP-Drp1 bound as the ratio of GFP-Drp1-WT:Drp1-S579D increases (**Figure 7A,B**), suggesting no effects of Drp1-S579D on Drp1-WT binding to filaments. Interestingly, there is a non-linear increase in actin-stimulated GTPase activity when GFP-Drp1-WT is titrated into Drp1-S579D, whereby no increase in GTPase activity is observed until 40% GFP-Drp1-WT is present (**Figure 7C**). A similar non-linear effect of Drp1-S579D occurs when un-tagged Drp1-WT is used (**Figure 7D**), indicating that the GFP-tag is not the source of the effect. This non-linear effect of Drp1-S579D on actin-activated Drp1-WT GTPase activity suggests that the S579D mutant slightly inhibits Drp1-WT activation when heteromerically bound to actin.

**Figure 7.**
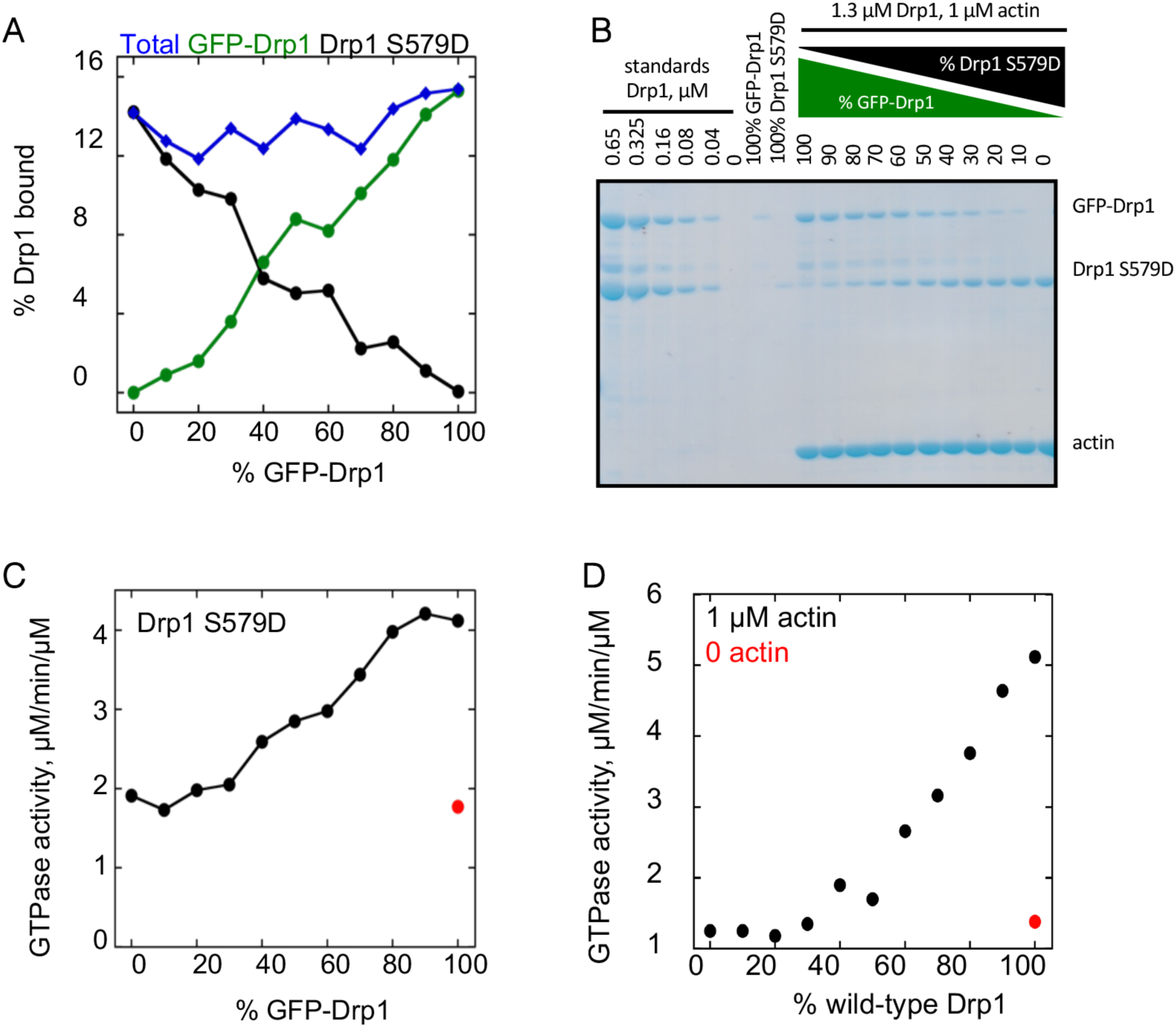
Drp1 phospho-mimetic mutants reduce the ability of actin filaments to activate wild-type Drp1. In all panels, the actin filament concentration is constant at 1 µM, and the total Drp1 concentration (combination of GFP-Drp1 and phospho-mimetic mutant Drp1 without GFP) is constant at 1.3 µM. **(A)** Graph of Drp1 binding to actin filaments with varying ratios of GFP-Drp1-WT:Drp1-S579D. **(B)** Corresponding Coomassie gels for binding assays quantified in panel A. **(C)** GTPase activity of Drp1 at varying ratios of GFP-Drp1-WT:Drp1-S579D in the presence of actin filaments. Red point represents GTPase activity of GFP-Drp1-WT in the absence of actin filaments. **(D)** GTPase activity of Drp1 (without the GFP tag) at varying ratios of Drp1-WT:Drp1-S579D in the presence of actin filaments. Red point represents GTPase activity of Drp1-WT in the absence of actin filaments.

We tested an additional mutant, Drp1-K38A, for effects on Drp1-WT activity under activating conditions. This mutant is widely used as a dominant-negative in cellular experiments. An equivalent mutant in dynamin 1 is also a dominant-negative in cells, and causes increased oligomerization of the wild-type protein ^39^. As expected, Drp1-K38A displays no GTP hydrolysis in the absence or presence of actin filaments (**Figure 8A**), while still binding actin filaments with similar properties to Drp1-WT (**Figure 8B,C**).

**Figure 8.**
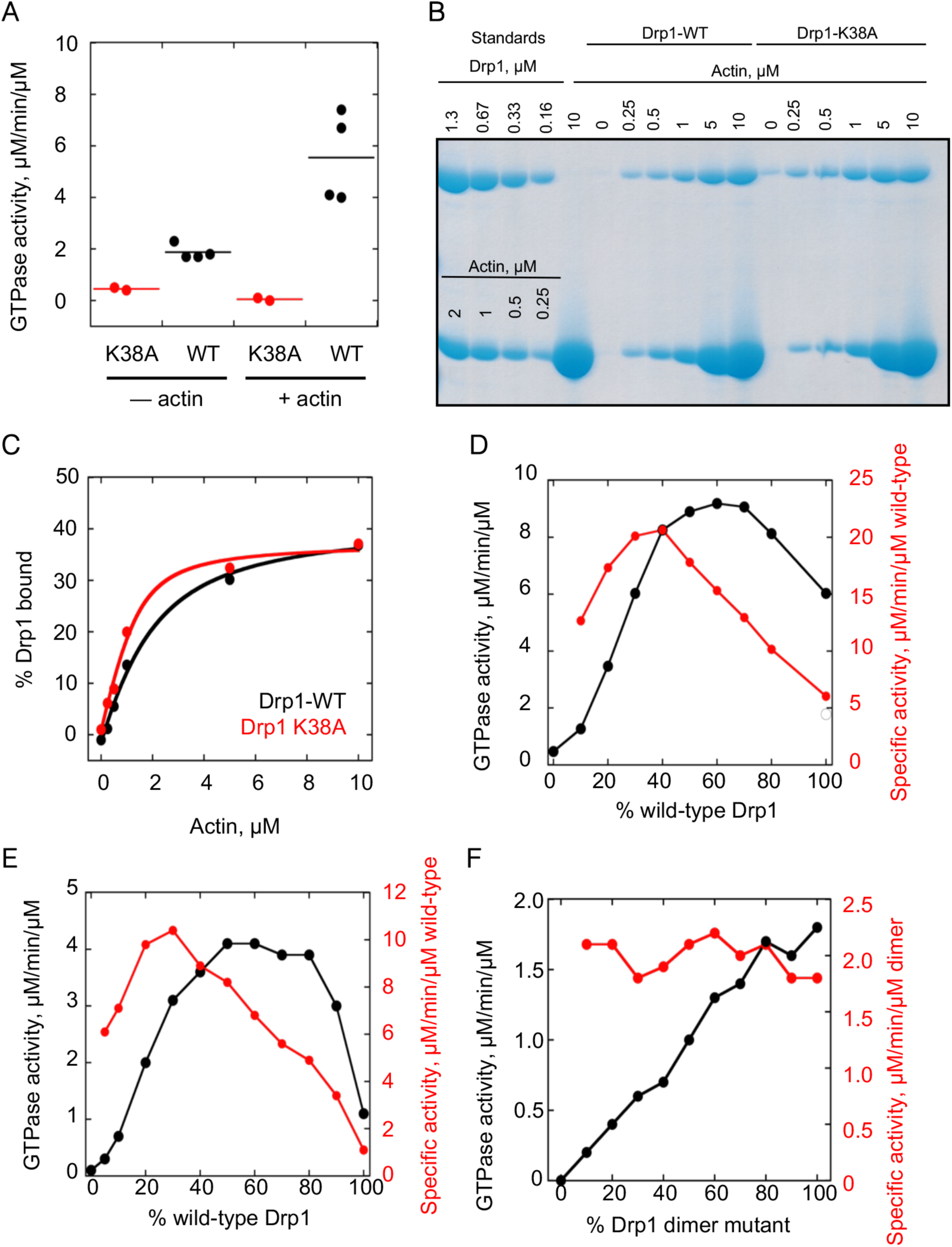
Drp1-K38A mutant stimulates the GTPase activity of Drp1-WT. (A) GTPase activity of Drp1-WT or Drp1-K38A (1.3 µM) in the presence or absence of actin filaments (1 µM). **(B)** Coomassie-stained SDS-PAGE of Drp1/actin co-sedimentation at 65 mM NaCl, with 1.3 µM Drp1-WT or Drp1-K38A and varying actin. Standards of known µM amounts of Drp1-WT and actin on left, pellets from sedimentation assays on right. **(C)** Graph of % Drp1 in the pellet for Drp1-WT and Drp1-K38A as a function of actin concentration. **(D)** GTPase activity of Drp1 at varying ratios of Drp1-WT:Drp1-K38A (1.3 µM total Drp1) in the presence of actin filaments (1 µM). Black line denotes total Drp1 specific activity (factoring both WT and K38A mutant) while red line denotes specific activity of Drp1-WT. **(E)** GTPase activity of Drp1 at varying ratios of Drp1-WT:Drp1-K38A (1.3 µM total Drp1) in the absence of actin filaments. Black line and red lines as described in panel D. **(F)** GTPase activity of Drp1 at varying ratios of Drp1-DM:Drp1-K38A (1.3 µM total Drp1) in the presence of actin filaments (1 µM). Black line denotes total Drp1 specific activity (factoring both Drp1-DM and K38A mutant) while red line denotes specific activity of Drp1-DM. DM, dimer mutant.

The absence of all GTPase activity for Drp1-K38A allowed us to test its effect on the specific activity of Drp1-WT directly, when the two are bound to actin filaments. Interestingly, Drp1-K38A has a significant stimulatory effect on actin-activated GTPase activity for Drp1-WT in these assays. The maximum effect occurs at 30-40% Drp1-K38A, where Drp1-WT specific activity is over 3-fold higher than for 100% Drp1-WT (**Figure 8D**). We also compared GTPase activity of varying ratios of WT and K38A mutants with no actin present. In this case, K38A has an even more dramatic effect, with an 80:20 ratio of K38A:WT having 10-fold higher specific activity than 100% Drp1 (**Figure 8E**). A similar effect has previously been observed for dynamin-1 ^39^.

To verify that the stimulatory effect of Drp1-K38A on Drp1-WT is due to hetero-oligomerization, we utilized the Drp1 dimer mutant (Drp1-DM) in place of Drp1-WT. Drp1-DM has similar basal GTPase activity to Drp1-WT ^2^, but remains dimeric at all concentrations tested and does not form a sedimentable species in the presence of GMP-PCP ^9^. As the percentage of Drp1-DM increases relative to Drp1-K38A, there is a linear increase in GTPase activity and no increase in specific activity (**Figure 8F**). We conclude that the stimulatory effect of Drp1-K38A on Drp1-WT GTPase activity is due to hetero-oligomerization effects.

These results suggest that Drp1-S579D reduces the activity of Drp1-WT when both are bound to actin filaments. In contrast, the GTPase activity of Drp1-WT is increased by the proximity of the catalytically inactive Drp1-K38A, either when bound to actin filaments or free in solution.

## Discussion

In this study, we have tested the effects of two phospho-mimetic mutants on Drp1 activity. The S579D mutant is widely thought to be stimulatory to Drp1 activity in cells ^16,18–25^, while the S600D mutant has been shown to be inhibitory in most cellular studies ^26,27,30,32^, but stimulatory in others ^28,29,31,33^. Both phospho-mimetic mutants display decreased oligomerization in a GTP-free state, but oligomerize in the presence of GTP. Both phospho-mimetic mutants display low GTPase stimulation by actin filaments, and reduced stimulation by cardiolipin or the Drp1 receptors Mff and MiD49. Drp1 phosphorylated *in vitro* on S579 displays similar attributes, with its cardiolipin-stimulated GTPase activity being even less than for its corresponding phospho-mimetic. When mixed with Drp1-WT on actin filamentsDrp1-S579D decreases the actin-stimulated activity of Drp1-WT.

The most striking aspect of these results is that phosphorylation at the S579 site, which has been associated with increased Drp1 activity in cellular studies, causes a decrease in activator-stimulated GTP hydrolysis, a fundamental read-out of biochemical Drp1 activity, GTP hydrolysis. One possible reason for this apparent contradiction is that the Drp1 isoforms used here: isoform 3 which contains none of the alternately spliced exons and isoform 6 which contains all three exons (**Figure 1**), do not display activation by S579 phosphorylation while other isoforms do. In the most comprehensive assessment to-date of Drp1 isoform abundance at the protein level ^16^, most cell and tissue types contain a mix of isoforms, with isoform 3 being the most abundant in some (but not all) cell lines. Nonetheless, other isoforms represent a significant proportion of total Drp1. The abundance of isoform 6 is unclear, but in commonly used cell lines such as HeLa is was not detected compared with four other isoforms ^16^. Considering that two alternatively spliced exons lie near the S579 site, additional studies are needed to test isoform-specific effects of phosphorylation.

Another possibility is that the positive effect of S579 phosphorylation depends on additional factors. For instance, S579 phosphorylation might result in recruitment of a factor that is stimulatory to Drp1 oligomerization, GTPase activity, or ability to constrict membranes. It is unclear what such a factor might be, but one possibility is a member of the nucleotide diphosphate kinase (NDPK) family that can catalyze GTP synthesis from GDP and ATP. Cytosolic NDPK family members NME1 and NME2 have been shown to increase activity of dynamin1 and dynamin2, while mitochondrial NME4 increases activity of the mitochondrial inner membrane dynamin Opa1 under conditions of low GTP ^40^. Physical interaction between these NDPKs and their respective dynamin proteins might raise the local concentration of GTP, increasing GTPase activity. There is evidence that another NDPK, NME3, enriches on peroxisomes and mitochondria, and might work with Drp1 in a similar manner ^41,42^.

Our results also provide information as to Drp1’s interaction with actin filaments. Both phospho-mimetic mutations, as well as S579 *in vitro* phosphorylation, eliminate actin-activated GTPase activity. This effect suggests that the Variable Domain is involved in the actin interaction. Interestingly, actin still seems to synergize with Mff for activation of the phospho-mimetics, suggesting that the important feature for this synergy is actin binding, not the activation that actin alone causes for Drp1.

It should be noted that, while the phospho-mimetic S579D mutant largely displays similar properties to S579-phosphorylated Drp1, the effects of phosphorylation are generally more pronounced. In particular, activation by cardiolipin-containing vesicles, as well as the synergy between actin and Mff, is practically eliminated for Drp1-phosphoS579, whereas it is partially inhibited for Drp1-S579D.

Finally, the inhibitory effect of Drp1-S579D on WT Drp1 GTPase activity is likely due to the decreased oligomerization of the mutant, which decreases the interactions between G domains necessary for GTPase activation. In contrast, the K38A mutation might activate WT Drp1 by stabilization of the oligomeric state. A similar effect of the equivalent mutation in dynamin-1, K44A, was described many years ago ^39^. These results emphasize the effect of oligomerization on Drp1 activity, with mechanisms that increase oligomerization being likely to increase GTPase activity.

**Experimental Procedures**

## METHODS

### Plasmids

For bacterial expression, full-length of human DRP1 isoform 3 (NP_055681.2, UniProt ID O00429-4), truncated human MFF isoform 4 (UniProt ID Q9GZY8-4) (MFF-ΔTM), and MiD49Δ1-124 (mouse amino acids 125-454, UniProt ID Q5NCS9) have been described previously ^7,9,43^. Quick Change mutagenesis was performed to make Drp1 mutants K38A, S579D, S600D, and Drp1 dimer mutant (Drp1-DM) ^2,9^.

### Protein Expression, Purification

DRP1 and its mutants were expressed and purified as previously described with modifications^9^. Briefly, DRP1 construct was expressed in One Shot BL21 Star (DE3) *Escherichia coli* in LB broth, induced by isopropyl-β-D-thiogalactoside (IPTG) at 16 °C for 16 hours when OD600 reached to 1.5. Cell pellets were resuspended in lysis buffer (100 mM Tris-Cl, pH 8.0, 500 mM NaCl, 1 mM dithiothreitol [DTT], 1 mM Ethylenediaminetetraacetic acid [EDTA], 2 µg/ml leupeptin, 10 µg/ml aprotinin, 2 µg/ml pepstatin A, 2 mM benzamidine, 1 µg/ml ALLN, and 1 µg/ml calpeptin) and lysed using a high-pressure homogenizer. The lysate was cleared by centrifugation at 40,000 rpm in Ti-45 rotor for 1 hour at 4°C. Avidin (20 µg/ml; PI-21128; Thermo Fisher Scientific, Waltham, MA) was added to the supernatant, and then was loaded onto Strep-Tactin Superflow resin (2-1206-025; IBA, Göttingen, Germany) by gravity flow. The column was washed with 20 column volumes (CV) of lysis buffer without protease inhibitors. To elute DRP1, 0.01 mg/ml HRV3C protease in lysis buffer without protease inhibitors was added for 16 hours at 4°C. The Strep-Tactin Superflow eluate was further purified by size exclusion chromatography on Superdex200 with DRP1-S200 buffer (20 mM HEPES pH 7.5, 150 mM KCl, 2 mM MgCl2, 1 mM DTT, 0.5 mM EGTA), spin concentrated, frozen in liquid nitrogen, and stored at −80 °C.

MFF-ΔTM was expressed in Rosetta^TM^2 BL21-(DE3) *Escherichia coli* (71400; EMD Millipore Corporation, Burlington, MA) in LB broth, induced by 1M IPTG at 30 °C for 4 h when OD600 reached to 1.5. Cell pellets were resuspended in lysis buffer (50 mM Tris-HCl, pH 7.5, 500 mM NaCl, 20 mM imidazole, pH 7.5, 1 mM DTT, 1 mM EDTA, 2 µg/ml leupeptin, 10 µg/ml aprotinin, 2 µg/ml pepstatin A, 2 mM benzamidine, 1 µg/ml ALLN, and 1 µg/ml calpeptin) and lysed using M-110 microfluidizer processor. The lysate was cleared by centrifugation at 40, 000 rpm in Ti45 for 40 minutes at 4°C, the supernatant was saved. Affinity capture was performed using FPLC and a HiTrap IMAC column (17-5248-01, GE Healthcare, Chicago, IL) equilibrated with IMAC-A buffer (50 mM Tris-HCl pH 7.5, 0.1 M NaCl, 20 mM imidazole). Cleared lysate was loaded onto the column with a rate of 3 mL/min and washed to baseline with IMAC-A. MFF was eluted from the column with gradient step washes by IMAC-B buffer (50 mM Tris-HCl pH 7.5, 0.1 M NaCl, 500 mM imidazole): step1 10% IMAC-B for 5CV, step2 20% IMAC-B for 5CV, step3 100% for 5CV. Fractions from step3 were pooled and diluted 10-fold in ion exchange (IEX)-A buffer (50 mM Tris-HCl pH 7.5, 1 mM DTT). Diluted fractions were loaded onto a HiTrap Q anion exchange column (54816, EMD Millipore Corporation, Burlington, MA). The column was washed to baseline with IEX-A and MFF was eluted by IEX-B buffer (50 mM Tris-HCl pH 7.5, 1 M NaCl, 1 mM DTT) with a step gradient: step1 10% 5CV, linear 10-50% 30CV followed by linear 50-100% 5CV. Peak MFF fractions were concentrated by reloading onto the HiTrap IMAC column and eluted with 100% IMAC-B step wash. MFF fractions were pooled and further purified by size exclusion chromatography on Superdex200 with S200 buffer (20 mM HEPES, pH 7.4; 2 mM MgCl2, 0.5 mM EGTA, 65 mM KCl, 1 mM DTT), spin concentrated (UFC903024, EMD Millipore Corporation, Burlington, MA), aliquots were frozen in liquid nitrogen, and stored at −80 °C.

MiD49 was expressed in One Shot BL21 Star (DE3) *Escherichia coli* (C6010-03; Life Technologies, Carlsbad, CA) in LB broth, induced by isopropyl-β-D-thiogalactoside (IPTG) at 16 °C for 16 h when OD600 reached 1.5. Cell pellets were resuspended in MiD lysis buffer (25 mM 4-(2-hydroxyethyl)-1-piperazineethanesulfonic acid [Hepes], pH 7.4, 500 mM NaCl, 1 mM dithiothreitol [DTT], 2 µg/ml leupeptin, 10 µg/ml aprotinin, 2 µg/ml pepstatin A, 2 mM benzamidine, 1 µg/ml calpain inhibitor I [ALLN], and 1 µg/ml calpeptin) and lysed using a high-pressure homogenizer (M-110L Microfluidizer Processor; Microfluidics, Newton, MA). The lysate was cleared by centrifugation at 40,000 rpm (type 45 Ti rotor; Beckman, Brea, CA) for 1 hour at 4°C and then was loaded onto Pierce™ Glutathione Agarose (16101; Thermofisher) by gravity flow. The column was washed with 20 column volumes (CV) of lysis buffer without protease inhibitors. To elute MiD49/51, 1 unit/µL thrombin (T4648, Sigma-Aldrich) or 0.01 mg/ml HRV3C protease in lysis buffer without protease inhibitors was added for 16 hours at 4°C. The protein eluate was captured by HiTrap IMAC column (17-5248-01, GE Healthcare, Chicago, IL) and eluted by IMAC-B buffer (50 mM Tris-HCl pH 7.5, 0.1 M NaCl, 500 mM imidazole). The His-trap protein eluate was further purified by size exclusion chromatography on Superdex200 (GE Biosciences, Piscataway, NJ) with S200 buffer (20 mM Hepes, pH 7.4, 65 mM KCl, 2 mM MgCl2, 1 mM DTT, 0.5 mM ethylene glycol tetraacetic acid [EGTA]), spin concentrated (UFC903024, EMD Millipore Corporation, Burlington, MA), frozen in liquid nitrogen, and stored at −80 °C.

Rabbit skeletal muscle actin was extracted from acetone powder as previously described ^44^, and further gel-filtered on Superdex 75 16/60 columns (GE Healthcare). Actin was stored in G buffer (2 mM Tris, pH 8.0, 0.5 mM DTT, 0.2 mM ATP, 0.1 mM CaCl2, and 0.01% NaN3) at 4°C.

### *In vitro* phosphorylation of Drp1 by ERK2

For *in vitro* phosphorylation assay, 30µM purified Drp1 or Drp1-S600D was incubated with 100 nM ERK2 kinase (Thermo Fisher PV3313) at 30 °C for 4hrs. Reaction was conducted in 25mM Hepes, pH 7.4, 150 mM KCl, 10 mM MgCl2, 2 mM DTT, 1 mM ethylene glycol tetraacetic acid [EGTA]), 1% Thesit. Phosphorylated Drp1 was further purified by size exclusion chromatography on Superdex200 with S200 buffer (20 mM HEPES, pH 7.4; 2 mM MgCl2, 0.5 mM EGTA, 65 mM KCl, 1 mM DTT), spin concentrated (UFC903024, EMD Millipore Corporation, Burlington, MA), aliquots were frozen in liquid nitrogen, and stored at −80 °C.

### Phospho-Drp1 analysis by mass spectroscopy

*In vitro* phosphorylated Drp1 were diluted in SDS-PAGE sample buffer and resolved by SDS-PAGE. Gels were stained with Coomassie blue staining. Bands were excised from the gel (2 µg Drp1) and analyzed for phosphorylated peptides by the Taplin mass spectrometry facility (Harvard Medical School).

### Actin preparation for biochemical assays

For high-speed pelleting assay, actin filaments were polymerized from 20 µM monomers for 3 h at 23 °C by addition of a 10x stock of polymerization buffer (200 mM HEPES, pH 7.4, 650 mM KCl, 10 mM MgCl2, 10 mM EGTA) to a final 1x concentration. For GTPase assay, actin monomers in G-buffer were incubated with AG1-X2 100–200 mesh anion exchange resin (Dowex; 1401241; Bio-Rad) at 4 °C for 5 min to remove ATP, followed by low-speed centrifugation. 20 µM actin filaments were polymerized as described before. To maintain ionic strength across all samples, an actin blank was prepared in parallel using G-buffer in place of actin monomers and used to dilute actin filaments as needed for each sample. DRP1 was diluted in MEHD buffer (20 mM HEPES, pH 7.4, 2 mM MgCl2, 0.5 mM EGTA, 1 mM DTT) to adjust the ionic strength to the same as S200 buffer before biochemical assays.

### Size exclusion Chromatography assays

Drp1 and Dpr1-phosS579 oligomeric distribution was determined by Superose 6 increase 10/300 GL SEC column (GE Biosciences) in S200 buffer (20 mM HEPES, pH7.4, 65 mM KCl, 2 mM MgCl2, 0.5 mM EGTA, 1 mM DTT). Protein at varying concentration was loaded onto the column in a total volume of 500 µL and gel-filtered with a flow rate of 0.4 mL/min.

### MiD49 oligomer preparation

For in making MiD49 oligomer, 100µM purified MiD49Δ1-124 were incubated with 500µM Palmitoyl-CoA (Sigma-Aldrich, P9716) at 37 ℃ for 1hrs. Reaction was conducted in with S200 buffer (20 mM HEPES, pH 7.4; 2 mM MgCl2, 0.5 mM EGTA, 65 mM KCl, 1 mM DTT). MiD49Δ1-124 mixture was further purified by size exclusion chromatography on Superdex200 with S200 buffer.

### Liposome preparation

All lipids were purchased from Avanti Polar Lipids (Alabaster, AL). Liposomes were prepared by extrusion through polycarbonate membranes of 250nm pore diameter. 0% Cardiolipin liposome contained 65% DOPC (850375P, Avanti Polar lipid) and 35% DOPE (850725, Avanti Polar lipid). 25% Cardiolipin liposome contained 40% DOPC, 35% DOPE, and 25% cardiolipin (840012C, Avanti Polar lipid).

### GTPase assay

DRP1 (0.75 µM) was mixed with indicated concentrations of MiD49, MFF and/or actin filaments in S200 buffer. Sample were incubated at 37 °C for 5 min. At this point, GTP was added to a final concentration of 500 µM to start the reaction at 37 °C. Reactions were quenched at designated time points by mixing 15 µL sample with 5 µL of 125 mM EDTA in a clear, flat-bottomed, 96-well plate (Greiner, Monroe, NC). Six time points were acquired for all conditions, either in a 12 min time range, or in a 45 min time range depending on reaction speed. Released phosphate was determined by addition of 150 µL of malachite green solution as previously described ^9^ Absorbance at 650 nm was measured 15 min after malachite green solution incubation. GTP hydrolysis rates were determined by plotting phosphate concentration as a function of time in the linear phase of the reaction.

### Velocity Analytical Ultracentrifugation

Analytical ultracentrifugation was conducted using a Beckman Proteomelab XL-A and an AN-60 rotor. For sedimentation velocity analytical ultracentrifugation, Drp1 and its mutants in S200 buffer (65/150 mM KCl, 1 mM MgCl2, 0.5 mM EGTA, 1 mM DTT, 20 mM HEPES, pH 7.4) was centrifuged at either 5,000 (for oligomer) or 35,000 (for monomer) rpm with monitoring at 280 nm. Data analyzed by Sedfit to determine sedimentation coefficient, frictional ratio, and apparent mass. Sedimentation coefficient reported is that of the major peak (at least 80% of the total analyzed mass) at OD280.

### High-speed pelleting assay

Interactions between DRP1 and actin were tested in the S200 buffer; DRP1 and actin were mixed as described and were incubated for 1 hr at room temperature in a 100 µl volume. After incubation, samples were centrifuged at 80,000 rpm for 20 min at 4°C in a TLA-100.1 rotor (Beckman). The supernatant was carefully removed. Pellets were washed three times with S200 buffer and then resuspended in 100 µl of SDS–PAGE sample buffer and resolved by SDS–PAGE (LC6025; Invitrogen, Carlsbad, CA). Gels were stained with Coomassie Brilliant Blue R-250 staining (1610400, Bio-Rad, Hercules, CA), and band intensity was analyzed using ImageJ software.

## Acknowledgements

We thank Arminja Kettenbach for initial examination of the phosphorylation state of bacterially-expressed Drp1, and Lando Ciripi for exposing himself to the outer surface. This work was supported by NIH R01 GM069818 and R35 GM112545 to HNH, and NIH P20 GM113132 to the BioMT COBRE.

## Supplemental Figures

**Supplemental Figure S1.**
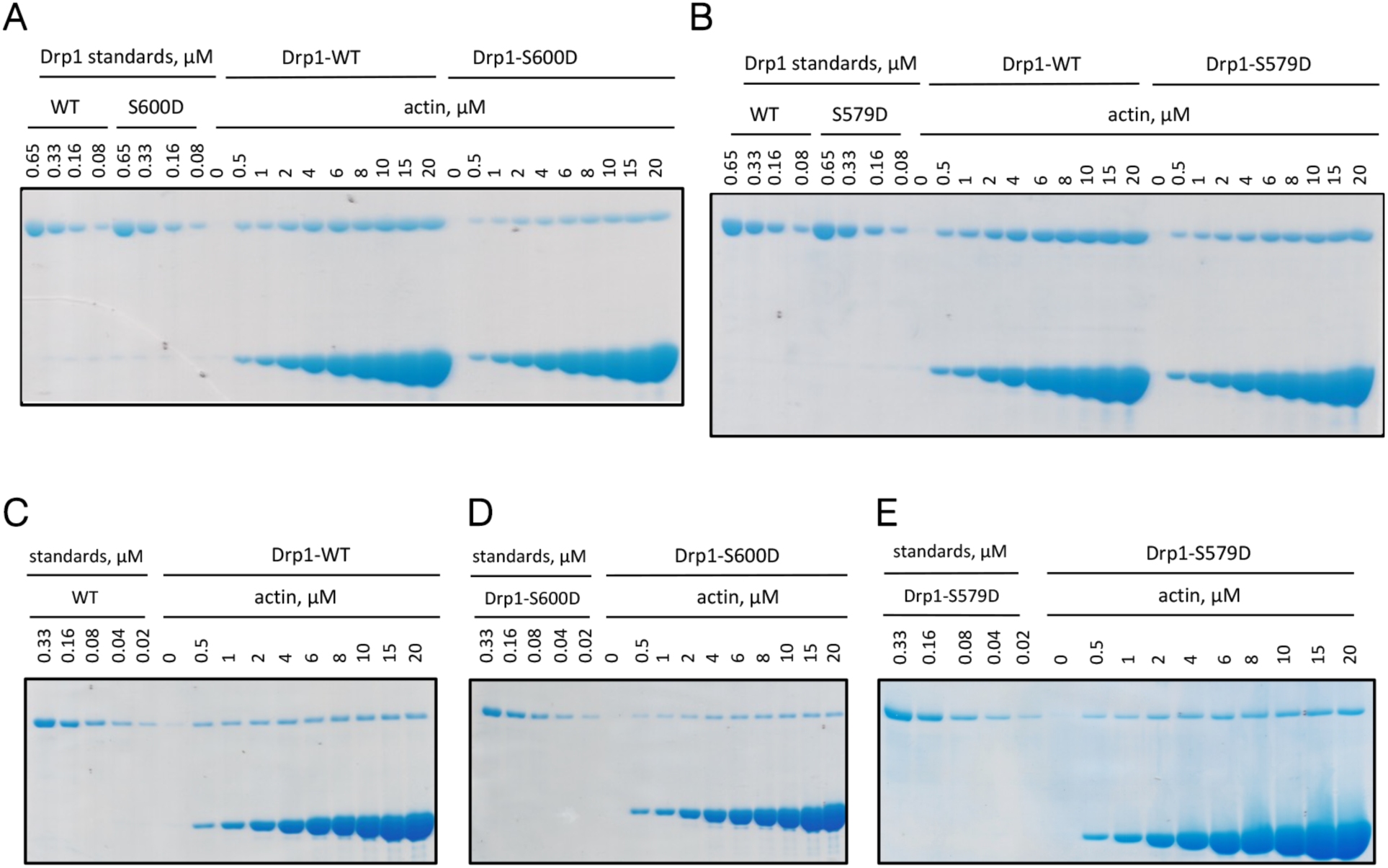
Drp1/actin co-sedimentation assays for phospho-mimetic mutants. A) and B) Coomassie-stained SDS-PAGE of Drp1/actin co-sedimentation assay at 65 mM NaCl similar to graph in Figure 3A. Standards of known µM amounts of Drp1 on left, pellets from sedimentation assays on right. A) Comparison of Drp1-WT with Drp1-S600D. B) Comparison of Drp1-WT with Drp1-S579D. C-E) Coomassie-stained SDS-PAGE of Drp1/actin co-sedimentation assay at 140 mM NaCl similar to graph shown in Figure 3B. Standards of known µM amounts of Drp1 on left, pellets on right. C) Drp1-WT. D) Drp1-S600D. E) Drp1-S579D. 1.3 µM Drp1 used in all assays.

**Supplemental Figure S2.**
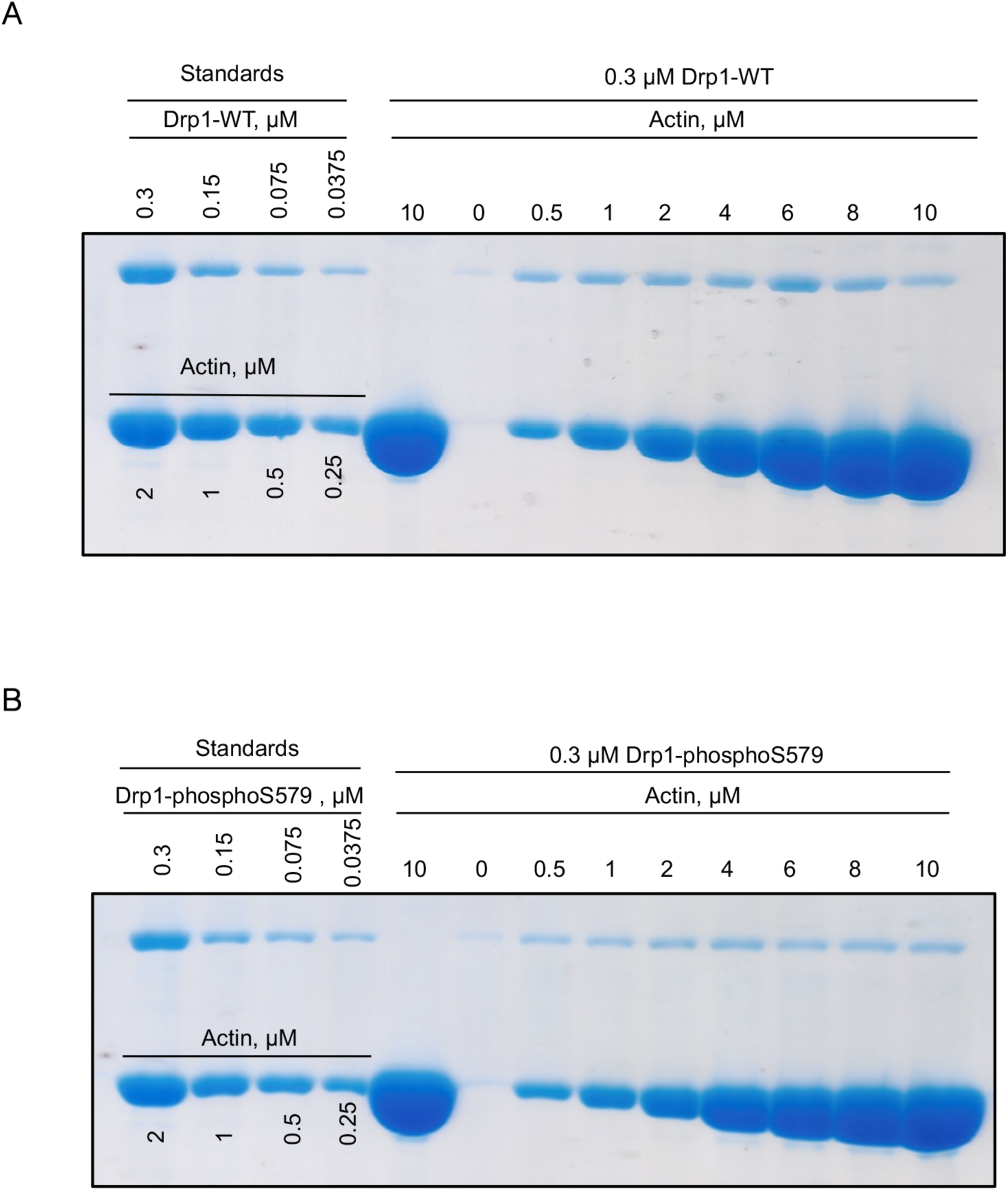
Drp1/actin co-sedimentation assays for ERK2-phosphorylated Drp1. A) and B) Coomassie-stained SDS-PAGE of Drp1/actin co-sedimentation assay at 65 mM NaCl similar to graph in Figure 4E. Standards of known µM amounts of Drp1 on left, pellets from sedimentation assays on right. A) Coomassie-stained SDS-PAGE of Drp1/actin co-sedimentation of Drp1-WT. B) Coomassie-stained SDS-PAGE of Drp1/actin co-sedimentation of Drp1-phosphoS579.

