## Supplemental figures for "Effects of phosphorylation on Drp1 activation by its receptors, actin, and cardiolipin"

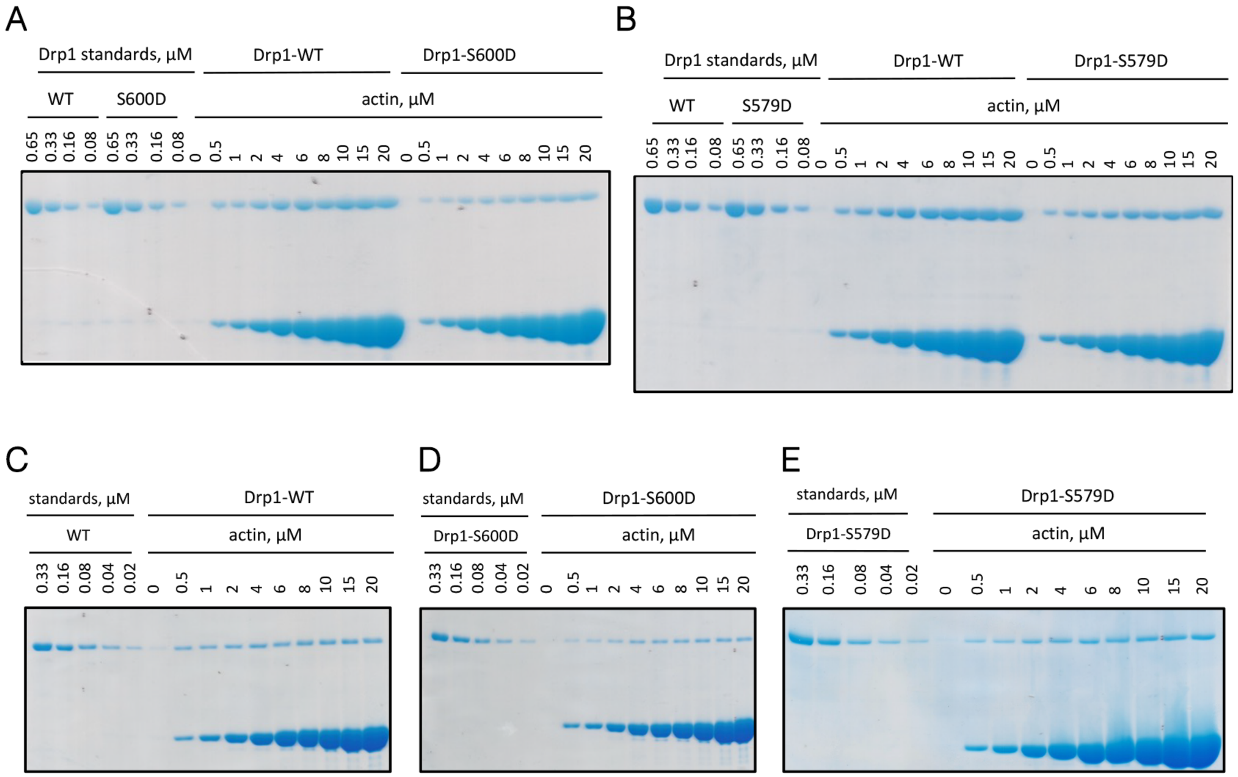


**Supplemental Figure S1. Drp1/actin co-sedimentation assays for phospho-mimetic mutants.** A) and B) Coomassie-stained SDS-PAGE of Drp1/actin co-sedimentation assay at 65 mM NaCl similar to graph in Figure 3A. Standards of known $\mu$M amounts of Drp1 on left, pellets from sedimentation assays on right. A) Comparison of Drp1-WT with Drp1-S600D. B) Comparison of Drp1-WT with Drp1-S579D. C-E) Coomassie-stained SDS-PAGE of Drp1/actin co-sedimentation assay at 140 mM NaCl similar to graph shown in Figure 3B. Standards of known $\mu$M amounts of Drp1 on left, pellets on right. C) Drp1-WT. D) Drp1-S600D. E) Drp1-S579D. 1.3 $\mu$M Drp1 used in all assays.

A

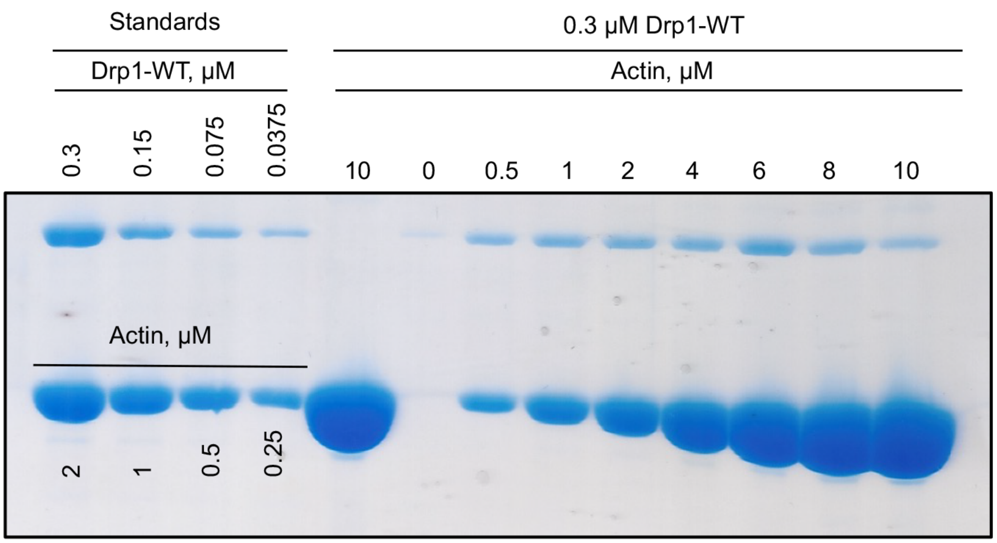


B

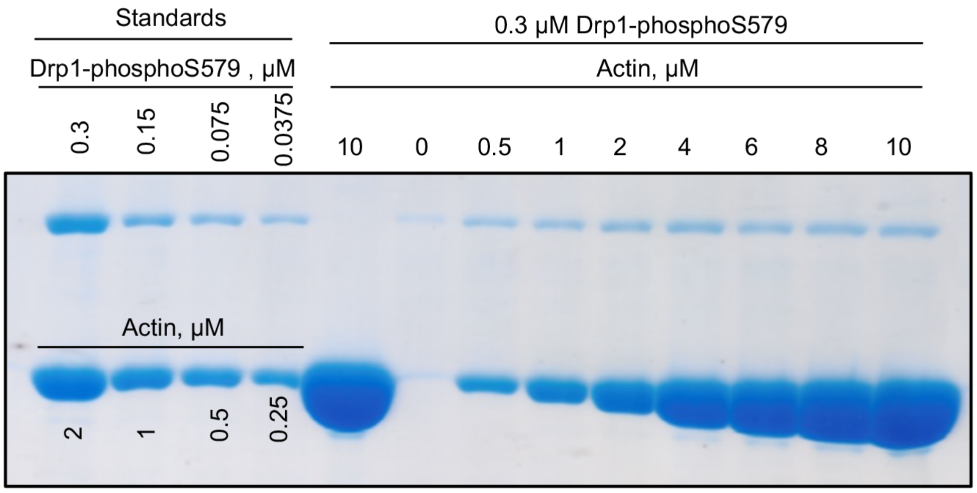


**Supplemental Figure S2. Drp1/actin co-sedimentation assays for ERK2-phosphorylated Drp1.** A) and B) Coomassie-stained SDS-PAGE of Drp1/actin co-sedimentation assay at 65 mM NaCl similar to graph in Figure 4E. Standards of known *μ*M amounts of Drp1 on left, pellets from sedimentation assays on right. A) Coomassie-stained SDS-PAGE of Drp1/actin co-sedimentation of Drp1-WT. B) Coomassie-stained SDS-PAGE of Drp1/actin co-sedimentation of Drp1-phosphoS579.
